# Experimental evolution reveals temporally opposing roles for Csu pili during *Acinetobacter baumannii* respiratory infection

**DOI:** 10.64898/2026.10.29.762983

**Authors:** Luca Cavinato, Clay D. Jackson-Litteken, Erin M Nawrocki, Neha Dubey, Chi Tai, Manon Janet-Maitre, Vaughn S Cooper, Christina L. Stallings, Gisela Di Venanzio, Mario F. Feldman

**Affiliations:** Department of Molecular Microbiology, Washington University School of Medicine in St. Louis, Missouri, USA; Department of Microbiology and Immunology, College of Medicine, University of Arkansas for Medical Sciences, Little Rock, Arkansas, USA; Department of Microbiology and Molecular Genetics, University of Pittsburgh, School of Medicine, Pittsburgh, Pennsylvania, USA

## Abstract

Multidrug-resistant (MDR) *Acinetobacter baumannii* is an opportunistic pathogen responsible for a wide range of infections, including ventilator-associated pneumonia, bloodstream infections, and urinary tract infections. Most studies investigating *A. baumannii* pulmonary pathogenesis rely on acute pneumonia models that use high bacterial inocula and do not recapitulate the prolonged infections observed in some patients. Recently, our laboratory developed a murine model of chronic pulmonary infection that enabled the identification of virulence factors specifically required for long-term lung persistence. To investigate how *A. baumannii* adapts during chronic pulmonary infection, we performed an experimental evolution study using this model. This approach revealed an unexpected role for the archaic chaperone-usher (Csu) pili in *A. baumannii* pathogenesis. At early time points, Csu expression reduced bacterial burdens in both acute and chronic pneumonia models, an effect associated with enhanced host inflammatory responses. In contrast, at later stages of chronic infection, Csu promoted bacterial persistence, potentially by facilitating adhesion to pulmonary epithelial cells. These opposing effects were conserved across two distinct *A. baumannii* clinical isolates. We also found that Csu contributes to bacterial colonization in a catheter-associated urinary tract infection (CAUTI) model in a strain-dependent manner. Together, these findings reveal an infection-stage-dependent role for Csu in *A. baumannii* pathogenesis, demonstrating that a bacterial surface structure can impair early colonization while promoting long-term persistence.

**Author summary:** Understanding how bacterial pathogens establish and maintain infections requires identifying the factors that promote their survival within the host. Currently, most experimental models of *Acinetobacter baumannii* infections focus on acute infections, potentially overlooking bacterial factors that are required for prolonged infections. In this study, we investigated how *A. baumannii*, a major cause of hospital-acquired infections, adapts to the lung environment during chronic infection. Using experimental evolution, we identified a novel role for Csu pili, surface structures traditionally associated with biofilm formation. Rather than promoting bacterial fitness throughout infection, Csu pili exhibited opposing effects depending on disease progression. Their expression was detrimental to bacterial fitness during early infection, a phenotype associated with increased host inflammatory responses, but became advantageous during prolonged infection, potentially by enhancing bacterial association with respiratory epithelial cells. Importantly, these opposing effects were observed in distinct clinical isolates, suggesting that this phenomenon is not restricted to a particular bacterial strain. Our findings highlight how the contribution of individual virulence factors can change over the course of infection and underscore the importance of considering infection duration when studying bacterial pathogenesis.

## Introduction

*Acinetobacter baumannii* is a Gram-negative opportunistic pathogen that has become a major global health concern because of its remarkable capacity to acquire antimicrobial resistance and persist in healthcare environments [1,2]. It causes a broad spectrum of infections, including hospital-acquired and ventilator-associated pneumonia, bloodstream infections, urinary tract infections (UTIs), and wound and soft-tissue infections, particularly in critically ill or immunocompromised patients [3–5]. The emergence of multidrug-resistant and carbapenem-resistant strains has substantially reduced therapeutic options, leading the World Health Organization to classify *A. baumannii* among the highest-priority bacterial pathogens for the development of new antimicrobial strategies [6,7].

*Acinetobacter baumannii* is a major cause of hospital-acquired pneumonia (HAP) [5,8] and is estimated to account for up to 10% of HAP cases in the United States [9,10]. Respiratory infections caused by *A. baumannii* are also associated with prolonged hospitalization, averaging approximately 30 days [11,12]. Despite its clinical significance, the bacterial factors that promote pulmonary persistence remain incompletely understood [5,8–10,13]. Although murine models of acute pneumonia have been instrumental in defining the mechanisms underlying *A. baumannii* pathogenesis [14–16], they are primarily designed to study early infection. These models typically involve intranasal or intratracheal inoculation with high bacterial doses (approximately 10⁷–10⁹ CFU), after which wild-type mice either clear the infection or succumb within 72 h, depending on the infecting strain. Consequently, experimental readouts are generally limited to the first 24–36 h of infection, restricting the identification of factors specifically required for long-term persistence. Acute models have established important roles for virulence determinants such as outer membrane protein A (OmpA), the secreted metalloprotease CpaA, capsular polysaccharide, and lipooligosaccharide (LOS) [17–23]. To overcome this limitation, we recently established a chronic pulmonary infection model using TLR4-defective C3H/HeJ mice and a sublethal inoculum of contemporary clinical *A. baumannii* isolates, enabling bacterial persistence in the lungs for up to 21 days [24]. Using this model, we identified the adhesin InvL as a determinant of long-term pulmonary persistence that is dispensable during acute pneumonia, demonstrating that factors required for chronic infection may differ from those driving acute disease [24].

These findings highlight the importance of studying infection beyond the acute phase and suggest that additional determinants of pulmonary persistence remain to be identified.

Chaperone-usher pili (CUP) are filamentous surface structures that mediate adhesion to host and abiotic surfaces [25]. *A. baumannii* encodes three major CUP systems, Abp1, Abp2, and Csu [5,26,27]. Abp1 and Abp2 are classical CUP pili that bind fibrinogen through the tip adhesins Abp1D and Abp2D and contribute to catheter-associated urinary tract infection (CAUTI) [5,28]. Csu is an archaic CUP system best known for promoting attachment to abiotic surfaces, including polystyrene and other materials used in medical devices [29,30]. In contrast to its established role in biofilm formation, the contribution of Csu to infection of host tissues, particularly the respiratory tract, remains poorly defined.

Here, we used experimental evolution in the chronic lung infection model to identify bacterial adaptations associated with repeated passage through the pulmonary environment. At passage 6, both independently passaged lineages of the respiratory isolate G636 (L1P6 and L2P6) had lost a ∼74-kb plasmid, p2_G636, and plasmid loss altered expression of the chromosomally encoded Csu pili. This observation led us to investigate the role of Csu during infection. We show that Csu has a temporally opposing role in the respiratory tract: its expression is detrimental during the early phase of infection, when it enhances inflammatory signaling, but promotes persistence at later time points, likely in part through its contribution to epithelial association. These findings demonstrate how chronic infection models can reveal virulence phenotypes that are not apparent, and may even appear reversed, in conventional acute models.

## Results

### Serial passage through the chronic lung model drives recurrent genetic changes in *A. baumannii*

We have previously demonstrated that the chronic lung infection model is a useful tool for studying multiple aspects of *A. baumannii* infection [24]. Because this model revealed virulence determinants that are dispensable during acute infection, we reasoned that serial passage through the chronic pulmonary environment might select for genetic changes that expose additional traits involved in persistence. We therefore performed a pilot experimental evolution study to examine the adaptation of *A. baumannii* to the chronic lung environment (Fig 1A and Materials and Methods).

**Figure 1.**
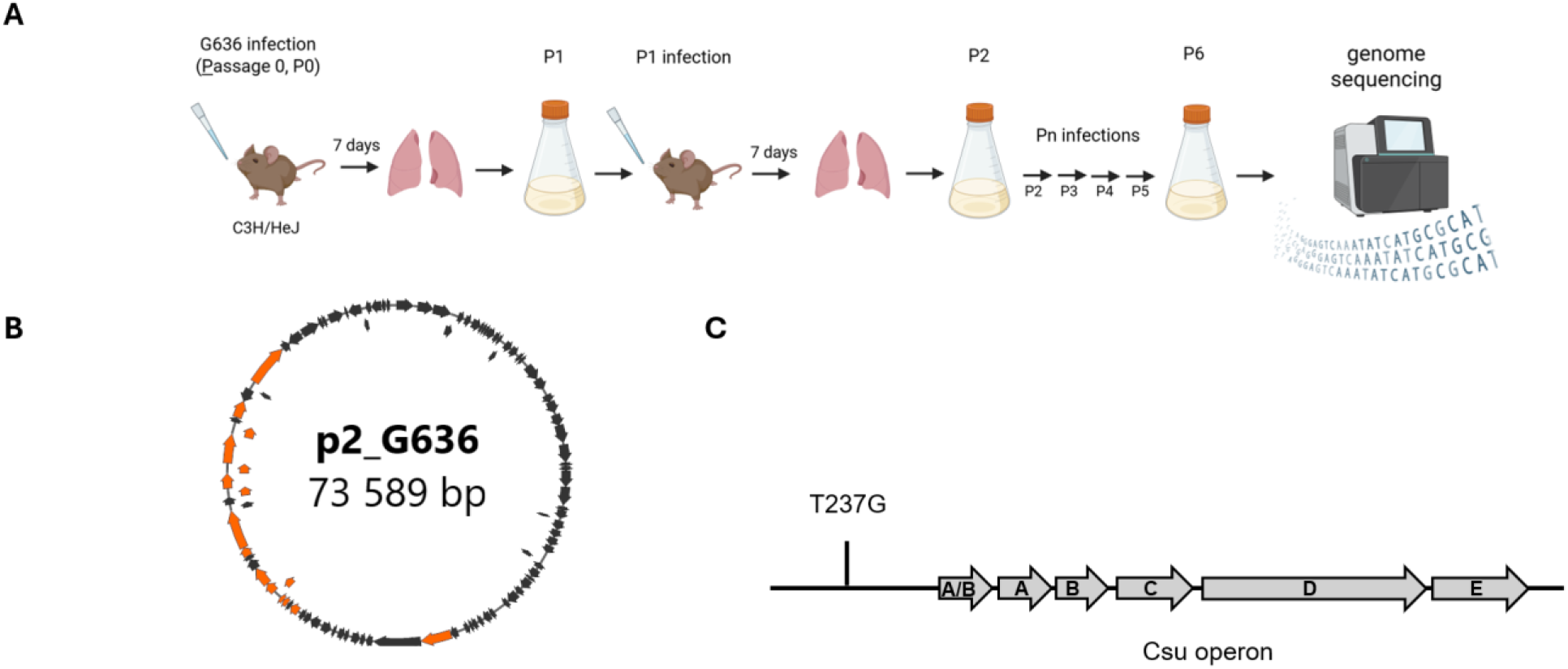
Experimental evolution of *A. baumannii* in the chronic lung infection model. A) Schematic representation of the six-passage evolution protocol. For each infection cycle, C3H/HeJ (*tlr4^-/-^*) mice were intranasally infected, and infection was allowed to persist for 1 week (see Materials and Methods). At each round, mice were euthanized, lungs collected, homogenized and cultured until they reached OD_600_ 0.5-0.8. Mid-log grown cultures were used as inoculum for the subsequent round of infection. At passage 6, single colonies were isolated from the population and whole-genome sequenced. B) Cartoon depicting p2_G636 plasmid. Orange arrows represent T4SS machinery; C) Schematic representation of SNP that occurred in lineage 2 upstream of the *csu* operon.

We selected the respiratory isolate G636 for these experiments as its persistence kinetics in the chronic lung model have been previously characterized [24]. We initially infected two C3H/HeJ *tlr4*-/- mice with the parental G636 isolate (P0) to establish two independent lineages, L1 and L2. After one week, mice were sacrificed, the lungs were homogenized, and bacteria recovered from each lung homogenate were grown in LB to mid-log phase (P1) and used as inocula for the subsequent round of infection (Fig 1A). After six passages, two independently evolved G636 populations, L1P6 and L2P6, were obtained.

To identify genetic changes that arose during experimental evolution, we performed whole-genome sequencing (WGS) of both P6 populations and four individual colonies from each lineage and compared them with the parental P0 strain (Figs 1B and 1C). Although individual colonies accumulated several distinct single-nucleotide polymorphisms (SNPs) throughout the genome (Tables A and B in S1 Appendix), we identified recurrent genetic changes that emerged during serial passage. Most notably, both lineages lost the conjugative plasmid p2_G636, one of the three plasmids carried by the parental strain (NZ_CP174092.1). p2_G636 is an approximately 74-kb conjugative plasmid found in *A. baumannii* [31].

In addition, the L2 population acquired a T→G substitution in the chromosome, located 237 bp upstream of the operon encoding the archaic CUP pili Csu, which mediate attachment to abiotic surfaces and biofilm formation [26,29,30]. This mutation, hereafter referred to as -237T>G, was present in three of the four sequenced L2 colonies, indicating that it had become predominant within the population. BLAST analysis of sequences deposited in the NCBI database identified the same nucleotide variant in 28 *A. baumannii* strains and one *A. pittii* strain, indicating that - 237T>G is a naturally occurring variant present in multiple *Acinetobacter* isolates.

Thus, serial passages through the chronic lung infection model reproducibly selected loss of p2_G636 and, in one lineage, and an additional chromosomal change, -237T>G, adjacent to the *csu* operon.

### Plasmid p2_G636 controls the expression of Csu pili

Because -237T>G mapped immediately upstream of the *csu* operon, we examined Csu levels in the four individual colonies isolated from each P6 evolved population by immunoblotting (Fig 2). All L1 colonies, which lacked p2_G636, displayed a dramatic reduction in Csu levels (Fig 2A, left panel). In contrast, three of the four L2 colonies, which also lacked the plasmid but carried the - 237T>G mutation, showed restored Csu levels. The remaining colony, L2C1, which lacked p2_G636 but had not acquired -237T>G, displayed Csu repression comparable to that observed in the L1 colonies (Fig 2A, right panel). Thus, loss of p2_G636 was consistently associated with Csu repression, whereas acquisition of -237T>G restored Csu production in the plasmid free background.

**Figure 2.**
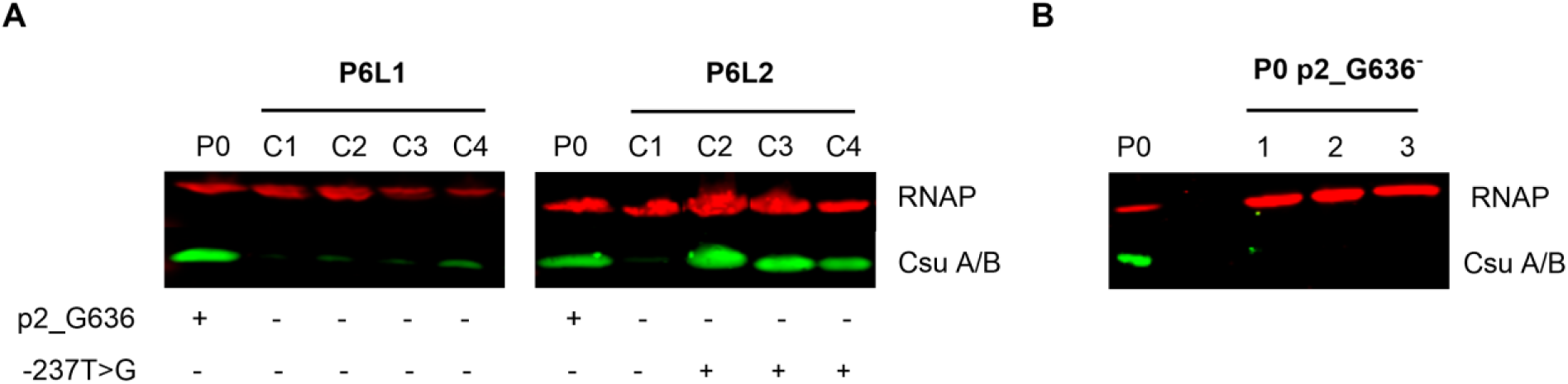
p2_G636 controls Csu levels. Western blot of whole-cell lysates. A) Single colonies (1-4) from Lineage 1 (L1, left panel) and Lineage 2 (L2, right panel) from passage 6 or P0 were visualized. B) P0 colonies cured of p2_G636. RNAP, RNA polymerase; Csu A/B, major csu pilin subunit.

To determine whether loss of p2_G636 was directly responsible for Csu repression, we took advantage of a recently described scar-free molecular tool for manipulating native plasmids in *A. baumannii* [32]. Briefly, the suicide vector pEMGT, carrying upstream and downstream homologous regions flanked by two recognition sites for the SceI endonuclease, was integrated into p2_G636. Subsequent expression of SceI targeted the plasmid and promoted its loss. As shown in Fig. 2B, curing p2_G636 from the parental P0 strain dramatically reduced Csu levels, confirming that the plasmid is associated with enhanced expression of the pili.

*A. baumannii* G636 also encodes two additional CUP pili, Abp1 and Abp2, whose expression was shown to be regulated by large conjugative plasmids (LCPs) in *A. baumannii* UPAB1 strain [5]. However, we could not detect Abp1 or Abp2 expression by immunoblotting in either P0 or the evolved strains under the conditions tested (S1 Fig). Together, these findings reveal a genetic interaction between p2_G636 and the chromosomal *csu* locus, whereby loss of the plasmid represses Csu protein levels and the -237T>G mutation seems to restore it.

### Csu pili have opposing effects during early and long-term lung infection

The major phenotypic difference associated with the recurrent genetic changes identified during experimental evolution was the differential expression of Csu between the two evolved lineages. To determine the consequences of this difference during infection and assess the contribution of p2_G636 to virulence, we infected C3H/HeJ *tlr4^-/-^* mice with P0, L1C3, or L2C2. L1C3 and L2C2 had both lost p2_G636, but only L2C2 carried the -237T>G mutation and expressed Csu in vitro. Hereafter, L1C3 and L2C2 are referred to as P6 Csu⁻ and P6 Csu⁺, respectively.

At 1 day post infection (dpi), mice infected with P6 Csu^-^ exhibited significantly higher bacterial burdens than those infected with P0 (Fig 3A). Bacterial burdens were also higher for P6 Csu^-^ than for P6 Csu^+^, although this difference did not reach statistical significance. By 7 dpi, bacterial burdens were comparable among all three groups (S2A Fig). Strikingly, by 12 dpi, the early phenotype was reversed: both P0 and P6 Csu^+^ exhibited significantly higher bacterial burdens than P6 Csu^-^ (Fig 3B).

**Figure 3.**
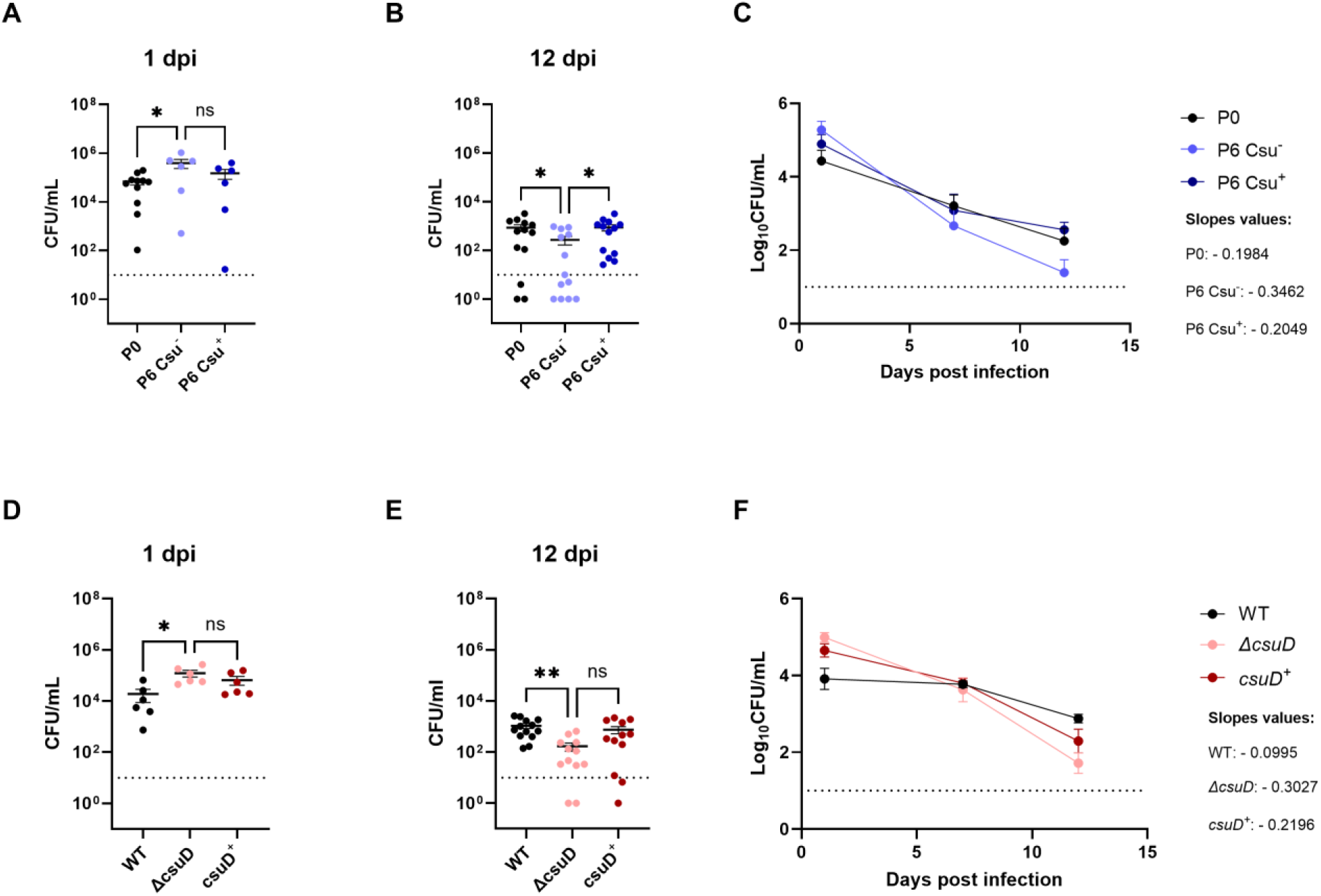
Csu pili have opposing effects during early and chronic pulmonary infection. A-F) C3H/HeJ mice were intranasally infected with the indicated strains. Bacterial burdens at 1 (A and D) and 12 (B and E) dpi are plotted. C and F) CFU from 1, 7, and 12 dpi were used for constructing bacterial killing curves. Linear regression model was used to calculate slopes of the curves. For the dot plots, each data point represents an individual mouse, the horizontal line represents the mean, and the SEM is indicated by error bars. Kruskal-Wallis test with Dunn’s multiple comparison was used. *, p<0.05; **, p<0.01.

Consistent with these observations, linear regression analysis of the bacterial clearance curves revealed a significant difference among their slopes (p = 0.0291), with P6 Csu^-^displaying the greatest absolute slope and therefore the fastest rate of clearance from the lungs (Fig 3C). These findings indicate that loss of Csu expression provides a transient advantage during the early stages of infection but compromises long-term persistence. Conversely, Csu expression appears detrimental initially following infection but becomes important for sustained infection of the lung at later time points.

A Δ*csuD* mutant, lacking the usher required for assembly and surface localization of Csu pili, exhibited a phenotype similar to that of P6 Csu^-^ in the chronic infection model, supporting the conclusion that the differences observed between the evolved strains were attributable to Csu pili. At 1 dpi, the Δ*csuD* mutant exhibited an approximately 1-log higher bacterial burden than the WT and *csuD*^+^ strains (Fig 3D). In contrast, at 12 dpi, the WT exhibited an approximately 1-log higher bacterial burden than the Δ*csuD* mutant (Fig 3E). This phenotype was partially complemented in the *csuD*^+^ strain, although the difference did not reach statistical significance. Consistent with our previous observations, analysis of the bacterial clearance curves further supported a role for Csu pili in long-term persistence in the lung. The slopes of the clearance curves were significantly different (p = 0.0001), with the Δ*csuD* mutant displaying the greatest absolute slope, indicative of more rapid clearance (Fig 3F). Together, these results confirm that loss of functional Csu pili provides an advantage early during infection but impairs the ability of *A. baumannii* to persist in the lung at later time points.

### Csu pili promote early inflammatory signaling

We next asked whether Csu pili contribute to innate immune recognition and thereby promote the enhanced bacterial clearance observed at 1 dpi in Csu-expressing strains. Using J774-Dual cells, which carry an NF-κB-inducible secreted alkaline phosphatase reporter, we found that the Δ*csuD* mutant induced significantly lower NF-κB activation than the WT and *csuD*^+^ strains (Fig 4A). This difference was not explained by differences in bacterial uptake (S3 Fig), indicating that functional Csu pili enhance inflammatory signaling independently of bacterial internalization.

**Figure 4.**
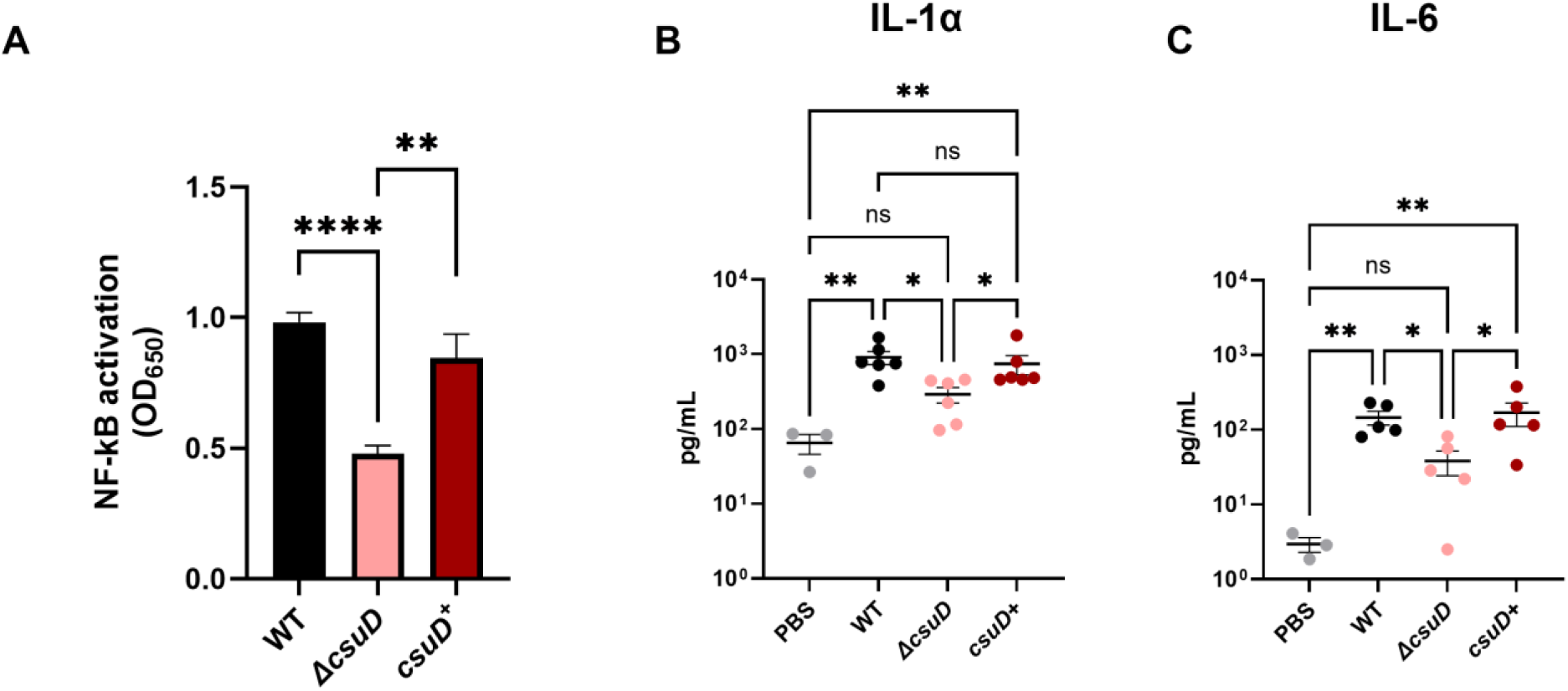
Csu pili contribute to early inflammatory responses. **A)** J774-Dual macrophages were infected for 2 hours at a MOI of 5. At the end of infection, supernatant was collected, centrifuged and analyzed for NF-κB activation. NF-kB activation is reported as OD_650_ values. B–C) Production of IL-1α (B) and IL-6 (C) was measured by flow cytometry in lung homogenates. PBS inoculated mice were used as negative control. Each data point represents an individual mouse, the horizontal line represents the mean, and the SEM is indicated by error bars. Kruskal-Wallis test with Dunn’s multiple comparison was used. *, p<0.05; **, p<0.01; ****, p<0.0001.

We next analyzed a panel of 13 cytokines in lung homogenates from infected mice (Table C in S1 Appendix). Although most cytokines were produced at similar levels at 1 dpi, IL-1α and IL-6 were significantly increased in infected lungs compared with PBS controls and were produced at significantly lower levels in mice infected with Δ*csuD* than in those infected with WT or *csuD*^+^ strains (Figs 4B and 4C). IL-1α functions as an alarmin released in response to tissue damage, whereas IL-6 is an important mediator of protective inflammatory responses [33,34].

At 12 dpi, concentrations of most cytokines were markedly reduced compared with 1 dpi. At this later time point, only IL-10, an immunomodulatory cytokine involved in limiting excessive inflammatory responses [35], differed significantly among groups. This difference may reflect differences in bacterial burden and in the magnitude of the earlier inflammatory response rather than a direct effect of Csu.

Despite these differences in cytokine production, we detected no significant differences in the abundance of alveolar macrophages or neutrophils in the airways at 1 dpi (S4, S5 Fig), indicating that the early detrimental effect associated with Csu expression is not accompanied by altered recruitment or abundance of these immune cell populations. Collectively, these findings indicate that Csu pili promote an early inflammatory response during lung infection, characterized by increased NF-κB activation and production of IL-1α and IL-6. Because these experiments were performed in *tlr4*-deficient mice, this response occurs independently of TLR4 and raises the possibility that Csu pili, either directly or indirectly, are sensed by the host innate immune system.

### Csu pili promote epithelial cell association at high bacterial density

Having established that Csu pili promote long-term persistence in the lung, we next sought to investigate the mechanism underlying this phenotype. Because we have previously demonstrated that adhesion is important for *A. baumannii* persistence in this model [24], we asked whether Csu contributes to bacterial association with epithelial cells. We assessed adhesion using two cell lines, A549 and 5637, derived from human lung and bladder epithelia, respectively (S6 Fig). At a low multiplicity of infection (MOI = 1), no differences in adhesion were observed. In contrast, at a higher MOI (MOI = 50), the *ΔcsuD* mutant displayed a clear adhesion defect in both cell lines compared with the WT and *csuD^+^* strains. Thus, Csu pili contribute to epithelial cell association under conditions of higher bacterial density, consistent with previous observations that Csu-mediated interactions are density dependent [38]. These findings suggest a potential mechanism by which Csu promotes long-term persistence, by enhancing and stabilizing bacterial association with epithelial surfaces. Recently, Malmo and colleagues showed that Csu promotes the formation of three-dimensional bacterial aggregates through interactions between adjacent cells [39]. Thus, Csu may enhance association with epithelial surfaces by promoting bacteria-bacteria interactions that stabilize bacterial communities at the epithelial surface.

### Csu expression is detrimental during acute pneumonia but promotes G636 urinary tract colonization

To determine whether the early disadvantage associated with Csu was also evident in a highly inflammatory setting, we examined the acute pneumonia model, a septic model in which C57BL/6 mount a strong inflammatory response to a high inoculation of bacteria [15,24]. At 24 h post infection, the Δ*csuD* mutant exhibited higher bacterial burdens than the WT and *csuD*^+^ strains, consistent with the phenotype observed at 1 dpi in the chronic model (Fig 5A). However, the difference was substantially greater in the acute model, with the Δ*csuD* mutant reaching approximately 3-log higher bacterial burdens in the lungs than the WT and complemented strains. This pronounced phenotype is consistent with the exaggerated inflammatory response elicited by the high inoculum in this model, which is strongly influenced by TLR4 [38,39]. Bacterial burdens in the spleen and kidney were also higher in mice infected with Δ*csuD* (S7 Fig), indicating that Csu expression limits bacteremia and dissemination during acute infection.

**Figure 5.**
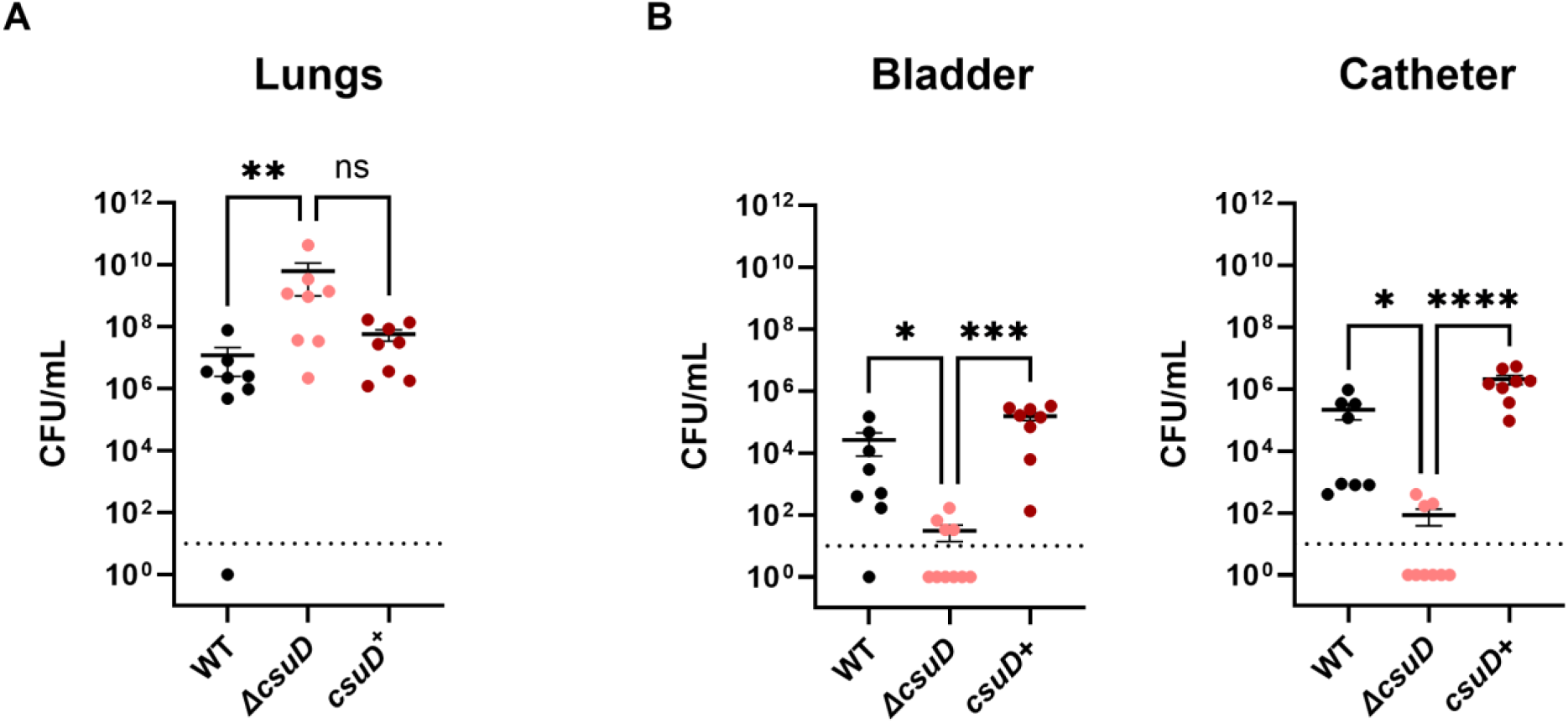
Csu expression is detrimental in the acute pneumonia model but promotes CAUTI. A) C57BL/6 mice were infected intranasally with the indicated strains; lungs were collected 24 hours post infection for assessment of bacterial burden. B) C57BL/6 mice were transurethrally infected after catheter insertion. Catheter (left) and bladder (right) were harvested 24 hours post infection. Each data point represents an individual mouse, the horizontal line represents the mean, and the SEM is indicated by error bars. Kruskal-Wallis test with Dunn’s multiple comparison was used. ns, not significant; *, p<0.05; **, p<0.01; ***, p<0.001; ****, p<0.0001.

CUP pili have been extensively studied in urinary tract infections, and we previously demonstrated the importance of *A. baumannii* CUP pili in a model of catheter-associated urinary tract infection (CAUTI) [5,28]. However, the contribution of Csu pili to UTI had not been examined. In the CAUTI model, the Δ*csuD* mutant was strongly attenuated, with most catheters and bladders showing little or no colonization at 24 h post infection compared with the WT (Fig 5B). Thus, in G636, Csu pili contribute to urinary tract colonization, consistent with their ability to enhance association with bladder epithelial cells and potentially with catheter-associated surfaces.

Together, these findings reinforce the opposing effects of Csu during infection. Csu pilus is detrimental during the early stages of lung infection, when it is associated with enhanced inflammatory signaling and bacterial clearance, but is beneficial in settings in which stable surface association and persistence are important.

### Csu has a conserved role in lung persistence across *A. baumannii* isolates

We next asked whether the in vivo phenotype observed in G636 was strain specific or more broadly conserved among *A. baumannii* isolates. To address this question, we generated a clean deletion of the entire *csu* operon (Δ*csu*) in the recent isolate Up398, which we have previously characterized for its ability to replicate within macrophages [40,41]. Similar to our observations with G636, the Δ*csu* mutant displayed an approximately 1-log higher bacterial burden in the lungs than the WT during acute infection (Fig 6A). This phenotype was also evident at systemic sites, with approximately 2-log higher bacterial burdens recovered from the spleens and kidneys of mice infected with Δ*csu* compared with WT (S8 Fig). Thus, loss of Csu also provides Up398 with an advantage during the early stages of infection.

**Figure 6.**
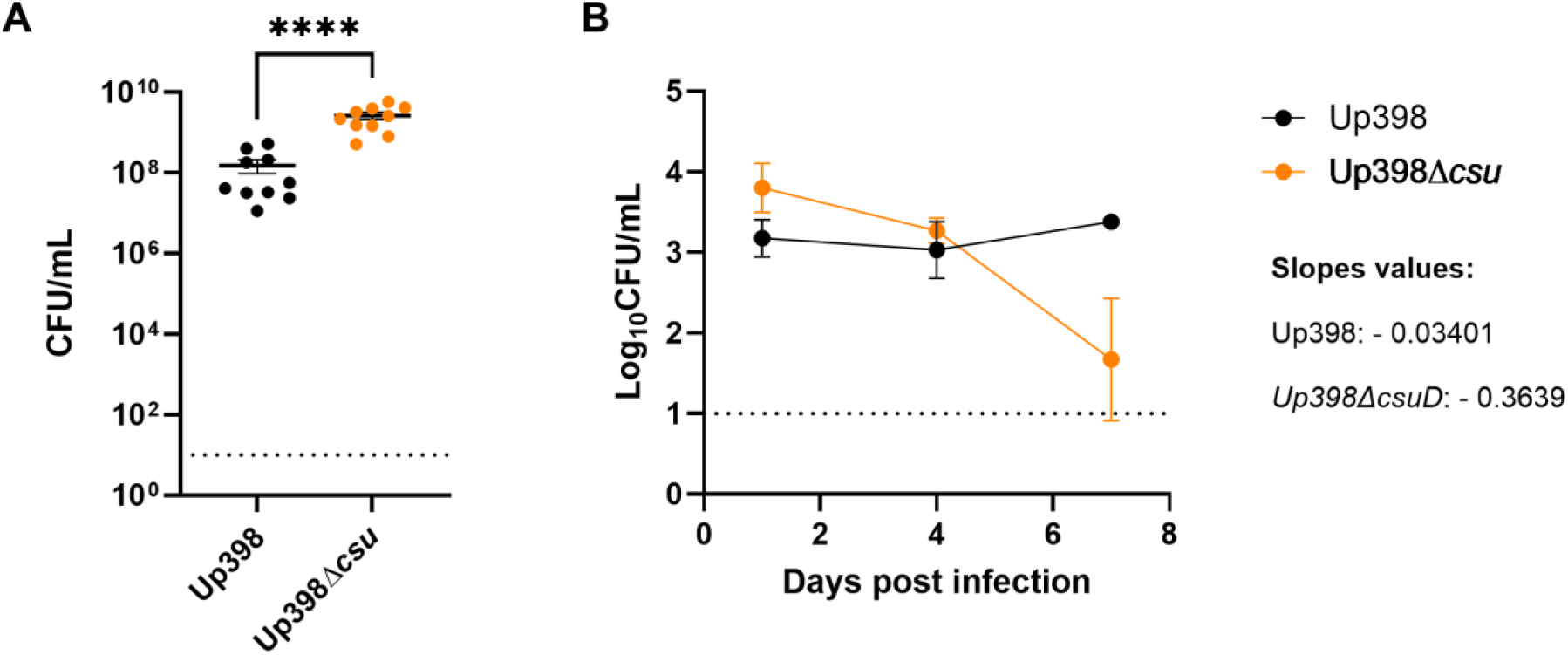
Csu contributes to lung persistence in the urinary isolate Up398. A) C57BL/6 mice were intranasally infected with the indicated strains. Lungs were collected 24 hours post infection for CFU count. Each data point represents an individual mouse; the horizontal line represents the mean. B) C3H/HeJ mice were intranasally infected with the indicated strains. Bacterial killing curves were constructed using CFU recovered at 1, 4 and 7 dpi. Linear regression model was used to calculate slope values of the curves. n = 3 for 1 and 7 dpi; n = 6 for 4 dpi. SEM is indicated by error bars. Kruskal-Wallis test with Dunn’s multiple comparison was used. ****, p < 0.0001.

We next examined the contribution of Csu to long-term lung persistence by infecting C3H/HeJ mice with a sublethal inoculum of WT or Δ*csu*. Consistent with the acute infection experiments, bacterial burdens were higher in the lungs of mice infected with Δ*csu* at 1 dpi (Fig 6B). However, as observed with the respiratory isolate G636, this phenotype was reversed at later stages of infection. By 7 dpi, the WT strain persisted at higher levels than Δ*csu*, and analysis of the bacterial clearance curves showed a more negative slope for the Δ*csu* mutant, indicating more rapid clearance from the lungs (Fig 6B). These findings confirm in a second *A. baumannii* isolate that Csu imposes a fitness cost early during infection but promotes persistence at later time points.

Notably, the kinetics of bacterial clearance differed between the two isolates. Whereas the persistence advantage associated with Csu became apparent by 7 dpi for Up398, the corresponding phenotype in G636 was most evident at 12 dpi (Figs 3F and 5A). These observations indicate that different *A. baumannii* isolates exhibit distinct persistence kinetics in the lung while retaining the same overall relationship between Csu presence and long-term persistence.

We next asked whether Csu also contributes to infection by Up398 in the CAUTI model (S9 Fig). In contrast to G636, no significant differences in bacterial burdens recovered from either the catheter or bladder were detected between WT and Δ*csu*. Thus, Csu is dispensable for Up398 fitness in the CAUTI model, suggesting that its contribution to urinary tract infection is strain-dependent or that other adhesins and virulence determinants can compensate for the absence of Csu in this isolate.

Together, these results extend our findings beyond G636 and demonstrate that the opposing effects of Csu during early- and long-term lung infection are conserved in distinct *A. baumannii* isolates. Csu expression is deleterious during the early stages of lung infection, when its absence promotes increased bacterial burden and dissemination, but becomes important for long-term persistence. In contrast, the contribution of Csu to CAUTI differs between G636 and Up398, highlighting the strain- and niche-dependent roles of these pili in *A. baumannii* pathogenesis.

## Discussion

In this study, we reveal an unexpected connection between plasmid carriage, Csu pili expression, and *Acinetobacter baumannii* pathogenesis in the lung. Loss of the p2_G636 plasmid resulted in strong repression of the chromosomally encoded *csu* operon, whereas a mutation upstream of *csu* restored Csu production in the plasmid-free background. Together, these findings identify p2_G636 as a previously unrecognized regulator of Csu expression and illustrate how changes in mobile genetic element content can reshape the expression of chromosomal determinants involved in host interactions.

Analysis of the resulting Csu phenotypes uncovered a striking time-dependent role for this pilus during respiratory infection. Early after infection, Csu expression imposed a substantial fitness cost. In both G636 and the unrelated clinical isolate Up398, bacteria lacking Csu reached higher burdens at 1 dpi. At later stages of infection, however, this relationship was reversed, and Csu-expressing bacteria displayed a survival advantage. Thus, Csu is neither simply beneficial nor detrimental during infection. Instead, its contribution changes over the course of disease: Csu expression is disadvantageous during the initial interaction with the host but promotes persistence once infection is established. These findings reveal that the contribution of a bacterial surface structure to pathogenesis can change substantially depending on the stage of infection.

Our results suggest that the early fitness cost associated with Csu expression may be linked to enhanced host inflammatory recognition. Csu expression increased NF-κB activation in macrophages and was associated with increased production of IL-1α and IL-6 in infected lungs. Importantly, this enhanced inflammatory response occurred despite lower bacterial burdens in mice infected with Csu-expressing bacteria, arguing that it does not simply reflect differences in bacterial numbers. IL-1α is an important alarmin associated with tissue damage, whereas IL-6 has broad pro-inflammatory functions [33,34]. These findings are consistent with a model in which Csu enhances host recognition during the early stages of infection, resulting in an inflammatory response that limits bacterial expansion. Previous work has similarly linked Csu expression and adhesion with increased IL-8 production and neutrophil responses [36], suggesting that enhanced inflammatory recognition may be an important and previously underappreciated consequence of Csu expression.

How Csu promotes this response remains unclear. Csu expression increased NF-κB activation in macrophages, raising the possibility that the pilus itself, or a Csu-dependent interaction with the cell surface, stimulates immune signaling. Alternatively, Csu-mediated adhesion to airway epithelial cells could enhance local inflammatory signaling and subsequently activate resident immune cells. Because the chronic infection model uses C3H/HeJ mice, which carry a loss-of-function mutation in TLR4, the Csu-dependent response is unlikely to be explained solely by canonical LPS-TLR4 signaling. Other pattern-recognition pathways, including TLR2 or intracellular receptors, could contribute [38,42–44]. Identifying the pathway responsible for Csu-dependent inflammatory signaling will require further investigation.

The early inflammatory cost associated with Csu contrasts with its contribution to bacterial survival later during infection. Csu has been studied primarily for its ability to mediate attachment to abiotic surfaces and biofilm formation [26,29,30], but increasing evidence supports a role in interactions with host cells [36,45]. Our previous work demonstrated that epithelial association becomes particularly important for *A. baumannii* survival during prolonged lung infection [24]. We therefore propose that Csu promotes late persistence by facilitating attachment to epithelial surfaces or the establishment of stable bacterial communities within the respiratory tract. Similar contributions of CUP pili to respiratory colonization have been observed in *Bordetella bronchiseptica* and *Burkholderia cenocepacia* [46,47]. Thus, the same surface structure that increases bacterial recognition early during infection may subsequently provide an advantage by promoting later retention within the host.

The context-dependent behavior of Csu may also help reconcile apparently conflicting observations for CUP pili in different infection models. Ahmed *et al.* reported reduced virulence of Csu-deficient *A. baumannii* in acute infection models [36], whereas we observed an early advantage associated with Csu loss. Differences in bacterial strain, host background, route of infection, inoculum, and experimental duration are all likely to influence the contribution of individual adhesins. A striking precedent comes from *Klebsiella pneumoniae* where deletion of *fimK* produces a hyperfimbriated phenotype that enhances biofilm formation and virulence during urinary tract infection but markedly impairs bacterial fitness in the lung [48,49]. We observed a similar dependence on infection context within our own study. Whereas G636 and Up398 displayed similar Csu-dependent phenotypes during respiratory infection, deletion of *csu* produced different outcomes in the CAUTI model. These observations emphasize that the contribution of CUP pili to pathogenesis is determined by bacterial background, anatomical niche, and stage of infection, and that increased expression of an adhesive structure cannot be assumed to uniformly enhance virulence. A limitation of the present study is that both strains examined lack detectable expression of the Abp1 and Abp2 CUP pili under the conditions tested. In isolates expressing these additional pili, functional redundancy may alter the consequences of Csu loss.

Beyond defining a role for Csu during respiratory infection, our findings reveal an unexpected relationship between plasmid carriage and regulation of a chromosomal surface structure. Plasmids are major drivers of *A. baumannii* biology through the accessory genes they encode, particularly determinants of antibiotic resistance. Our results show that their influence can also extend to regulation of chromosomal genes that directly affect interactions with the host. How p2_G636 controls the *csu* operon remains unknown and is beyond the scope of this study. However, the ability of the upstream -237T>G mutation to restore Csu expression in the absence of p2_G636 suggests that plasmid-dependent regulation intersects with the local regulatory circuitry controlling *csu*. Thus, acquisition or loss of a plasmid can have phenotypic consequences that extend well beyond the functions encoded by the plasmid itself.

Together, our findings reveal a dual and time-dependent role for Csu during respiratory infection. Csu expression carries an early cost associated with enhanced inflammatory recognition but provides an advantage during prolonged persistence. At the same time, we uncover an unexpected connection between plasmid carriage and regulation of this chromosomal surface structure. These findings highlight how the contribution of bacterial adhesins can change over the course of infection and reveal an additional mechanism by which mobile genetic elements can influence *A. baumannii* pathogenesis.

## Materials and Methods

### Plasmids, bacterial strains, and growth conditions

All plasmids and bacterial strains used in this study are listed in Table D in S1 Appendix. Bacteria were grown in Lennox broth or on Lennox agar at 37°C, either with shaking at 180 rpm or under static conditions, as indicated. When required, media were supplemented with chloramphenicol (10 µg/mL), apramycin (50 µg/mL for *A. baumannii* and 30 µg/mL for *Escherichia coli*), or potassium tellurite (50 µg/mL for *A. baumannii* and 25 µg/mL for *E. coli*).

### Genetic manipulation of bacteria

All primers used in this study are listed in Table E in S1 Appendix. Deletion mutants were generated by amplifying approximately 1,000 bp regions upstream and downstream of the target locus from genomic DNA using the indicated primer pairs. The suicide vector pEX18-Apr was amplified with vector-specific primers, and the overlapping fragments were assembled using NEBuilder HiFi DNA Assembly Master Mix (New England Biolabs). Recombinant plasmids were introduced into *E. coli* by electroporation and verified by sequencing (Plasmidsaurus). Mutational constructs were transferred into either G636 or Up398 (NCBI RefSeq assembly GCF_045256945.1 and GCF_050320275.1 respectively), and merodiploid strains were selected on Lennox agar containing chloramphenicol and apramycin. Double-crossover recombinants were obtained by counterselection on NaCl-free LB agar supplemented with 10% sucrose and incubated at room temperature for 48 h. Mutants were confirmed by PCR and whole-genome sequencing. Complementation was performed with pUC18T-miniTn7T-Apr::*csuD* and pUC18T-miniTn7T-Apr::*csu* using a four-parental conjugation strategy with pRK2013 and pTNS2 as helper plasmids. Complemented strains were selected with apramycin and chloramphenicol, and insertion at the mini-Tn7 site was confirmed by PCR.

Plasmid curing was performed as previously described with minor modifications [32]. Approximately 1,000 bp regions flanking a selected locus on p2_G636 were amplified and cloned into the suicide vector pEMGT. The plasmid was transferred into *A. baumannii* by four-parental mating, and integrants were selected on chloramphenicol and tellurite. The broad-host-range pSW plasmid was subsequently introduced using the same mating strategy. Loss of p2_G636 was confirmed by PCR and whole-genome sequencing.

### Murine models of infection

All animal experiments were approved by the Washington University Institutional Animal Care and Use Committee. Acute and chronic lung infections were performed as previously described [24] using 6- to 8-week-old female C57BL/6 and C3H/HeJ mice, respectively. Overnight bacterial cultures were diluted 1:200 in LB and grown for approximately 3 h to mid-log phase (OD_600_ 0.5-0.9), washed twice in PBS, and adjusted to the required inoculum. Mice were inoculated intranasally with approximately 10^8^ CFU for the acute model or 10^5^ CFU for the chronic model. At the indicated times post-infection, mice were euthanized and bacterial burdens in the lungs, spleen, and kidneys were determined by serial dilution and plating.

The murine CAUTI model was performed as previously described [5]. C57BL/6 mice were transurethrally implanted with a 4- to 5-mm segment of silicone tubing. Statically grown bacterial cultures were washed twice and resuspended in PBS, and mice were inoculated transurethrally with 50 µL containing approximately 2 × 10^8^ CFU. At 24 h post-infection, mice were euthanized and bacterial burdens in the bladder and on the catheter were quantified by serial dilution and plating.

### BAL collection and flow cytometry

After euthanizing the mouse, 1 ml cold PBS with 5mM EDTA was gently injected into the lung through the trachea, left for 5 seconds before retrieving the bronchoalveolar lavage (BAL) fluid. The flush was repeated for a total of 5 times and pooled. All samples were kept on ice. BAL cells were spun down at 500 x g for 5 min at 4°C. Total BAL cells were resuspended in semi-FACS buffer (1X PBS pH 7.4 supplemented with 2% FBS and 2 mM EDTA) in the presence of Fc receptor blocking antibody TruStain FcX™ PLUS (BioLegend, S17011E) and stained with fluorescent labelled tagged antibodies at a 1:200 dilution in a cocktail mix. Cell staining antibody cocktail included the following antibodies: CD45-BV510 (BioLegend, S18009F), Ly6G-BV421 (BioLegend, 1A8), CD11c-APC (BioLegend, N418), SiglecF-PerCP cy5.5 (BioLegend, S17007L), CD11b-AF700 (BioLegend, M1/70), and CD24-PE-Cy7 (BioLegend, 30-F1). Cells were stained at 4°C for 20 minutes and resuspended in semi-FACS buffer for flow cytometry. Flow cytometric analysis was performed on a Cytek Aurora and data was analyzed with FlowJo software (Tree Star).

### Experimental evolution in the chronic pneumonia model

In vivo serial passage of the respiratory isolate G636 was performed independently in two lineages, L1 and L2 (Fig. 1A). For passage 1, C3H/HeJ mice were intranasally inoculated with approximately 10^5^ CFU of mid-log-phase G636. After 7 days, mice were euthanized and the lungs were collected and homogenized. Lung homogenates were cultured in LB supplemented with chloramphenicol (10 µg/mL) to mid-log phase (OD_600_ approximately 0.5-0.9), and the recovered cultures were designated L1P1 and L2P1 and used as the inocula for the next round of infection. This procedure was repeated for six passages, generating the L1P6 and L2P6 populations.

### Whole-genome sequencing and comparative analysis

Genomic DNA was isolated from the parental G636 strain (P0), the evolved P6 L1 and P6 L2 populations, and four individual colonies from each P6 population. WGS libraries were prepared by SeqCoast (Portsmouth, NH) using the Illumina DNA Prep tagmentation kit with unique dual indexes and were sequenced as 2×150 bp paired-end reads on an Illumina NextSeq 2000. Raw reads were trimmed and assembled using Geneious Prime. To identify variants in evolved populations, sequencing reads were mapped to the G636 reference using breseq v0.39.0 (Deatherage and Barrick 2014) in polymorphism mode. Single-nucleotide variants and their frequencies in isolated colonies were identified in Geneious Prime and are reported in Supplementary Tables 1 and 2.

### Cell culture

A549 human lung epithelial cells (CCL-185) and 5637 human bladder epithelial cells (HTB-9) were cultured in high-glucose Dulbecco’s Modified Eagle Medium (DMEM) and RPMI 1640, respectively, supplemented with 10% heat-inactivated fetal bovine serum at 37°C and 5% CO2. J774-Dual reporter macrophages (InvivoGen) were cultured in high-glucose DMEM supplemented with 10% heat-inactivated fetal bovine serum at 37°C and 5% CO2.

### Cell infection assays

Adhesion assays were performed as previously described [50]. One day before infection, 2 × 10^5^ A549 cells or 3 × 10^5^ 5637 cells were seeded in 48-well plates. Overnight bacterial cultures were washed twice with PBS and normalized to OD_600_ = 1, then diluted in antibiotic-free medium to the indicated multiplicity of infection (MOI). Infected cells were centrifuged at 200 × g for 5 min and incubated for 1 h at 37°C and 5% CO2. Cells were then washed three times with PBS and lysed with 200 µL of 0.025% Triton X-100. Adherent bacteria were quantified by serial dilution and plating. Data were collected from at least three independent experiments.

J774-Dual reporter macrophages were infected as previously described with minor modifications [38]. Cells were seeded at 3 × 10^5^ cells per well one day before infection. Bacteria were prepared as described above and used at an MOI of 5. Cells were infected for 2 h at 37°C and 5% CO2. Supernatants were collected for reporter analysis, and cells were washed, lysed with 0.025% Triton X-100, and plated for bacterial enumeration.

### Western blotting

Whole-cell lysates were prepared from overnight static cultures. Bacteria were washed twice, pelleted, and resuspended in Laemmli sample buffer to a final OD_600_ of 10. Ten microliters of each sample were resolved on 12% SDS-PAGE gels and transferred to nitrocellulose membranes. Membranes were probed with polyclonal rabbit anti-CsuA/B (1:500) [29] and monoclonal mouse anti-RNA polymerase (1:3,500; BioLegend), followed by IRDye-conjugated anti-rabbit and anti-mouse secondary antibodies (1:5,000; LI-COR Biosciences). Blots were visualized using an Odyssey CLx imaging system.

### Cytokine measurements

Cytokines were quantified as previously described with minor modifications [24]. Lung homogenates were supplemented with protease inhibitor according to the manufacturer’s instructions, flash-frozen, and stored at -80°C. Samples were thawed and centrifuged at 10,000 × g for 12 min at 4°C to remove debris. Cytokine concentrations were measured using the LEGENDplex Mouse Inflammation Panel (13-plex; BioLegend) according to the manufacturer’s instructions and analyzed on a Cytek Aurora flow cytometer.

## Statistical analysis

Statistical analyses were performed using GraphPad Prism version 10. Data were assessed for normality, and parametric or nonparametric tests were selected as appropriate. Statistical significance was defined as P < 0.05.

## Data availability

Data generated in this study will be made available upon publication. Sequenced genomes of the *A. baumannii* evolved strains were deposited on NCBI (BioProject: PRJNA1545994). Prism software was used to display experimental data. The experimental workflows were made using bioRender (biorender.com).

## Acknowledgments

We thank Asya Smirnov for the training and use of Cytek Aurora flow cytometer. We thank Divman Gill and Dakota Hall for technical support, and the members of the Feldman lab for critical reading of the manuscript. We thank Lance Bottini for his diligent care of our instruments.

## Author contributions

Conceived and designed the experiments: L.C., G.DV. C.JL., and M.F.F. Performed the experiments: L.C., C.JL., G.DV., M.JM., N.D., C.T. Analyzed the data: L.C., E.M.N, V.S.C., N.D., C.L.S. Wrote the first draft of the manuscript: L.C. All authors edited the manuscript.

